# Comparison of Anorectal Function Measured using Wearable Digital Manometry and a High Resolution Manometry System

**DOI:** 10.1101/2020.01.24.917922

**Authors:** Ali Attari, William D. Chey, Jason R. Baker, James A. Ashton-Miller

**Affiliations:** Department of Mechanical Engineering, University of Michigan, Ann Arbor, MI, United States of America; Division of Gastroenterology and Hepatology, University of Michigan Hospitals and Health Centers, Ann Arbor, MI, United States of America; Department of Biomedical Engineering, University of Michigan, Ann Arbor, MI, United States of America; Institute of Gerontology, University of Michigan, Ann Arbor, MI, United States of America

**Keywords:** Digital manometry, high-resolution anorectal manometry, wearables, electromyography, dyssynergic defecation, chronic constipation, fecal incontinence

## Abstract

There is a need for a lower cost manometry system for assessing anorectal function in primary and secondary care settings. We developed an index finger-based system (termed “digital manometry”) and tested it in healthy volunteers, patients with chronic constipation, and fecal incontinence. Anorectal pressures were measured in 16 participants with the digital manometry system and a 23-channel high-resolution anorectal manometry system. The results were compared using a Bland-Altman analysis at rest as well as during maximum squeeze and simulated defecation maneuvers. Myoelectric activity of the puborectalis muscle was also quantified simultaneously using the digital manometry system. The limits of agreement between the two methods were −7.1 ± 25.7 mmHg for anal sphincter resting pressure, 0.4 ± 23.0 mmHg for the anal sphincter pressure change during simulated defecation, −37.6 ± 50.9 mmHg for rectal pressure changes during simulated defecation, and −20.6 ± 172.6 mmHg for anal sphincter pressure during the maximum squeeze maneuver. The change in the puborectalis myoelectric activity was proportional to the anal sphincter pressure increment during a maximum squeeze maneuver (slope = 0.6, R^2^ = 0.4). Digital manometry provided a similar evaluation of anorectal pressures and puborectalis myoelectric activity at an order of magnitude less cost than high-resolution manometry, and with a similar level of patient comfort. Digital Manometry provides a simple, inexpensive, point of service means of assessing anorectal function in patients with chronic constipation and fecal incontinence.

## 1. Introduction

High-resolution anorectal manometry (HR-ARM) has become the standard method for performing detailed evaluation of anorectal function [1]. HR-ARM provides information for the diagnosis of fecal incontinence (FI) and chronic constipation (CC) which affect up to 18% and 19%, respectively, of the adult population in North America [2–4]. More specifically, HR-ARM allows the identification of chronically constipated patients with dyssynergic defecation and FI patients with sphincter weakness, both of which are most effectively treated with physical therapy and biofeedback training rather than standard medical therapies like anti-diarrheals or laxatives.

A HR-ARM probe is comprised of an array of pressure sensors on a 4 mm diameter catheter and has been found to be more sensitive than conventional water perfused ARM (C-ARM) systems [5,6]. HR-ARM requires a relatively expensive system, an additional visit for the patient, and experienced staff to operate and interpret the results. Hence HR-ARM has become limited to tertiary care medical centers with the result that many patients are deprived of the opportunity to be properly diagnosed and triaged to appropriate therapy [7–9].

We hypothesized that a simpler and less costly system might be able to provide the most salient information provided by HR-ARM at the point of clinical service. Physicians have employed wearable devices to examine patients since Arthur Leared invented the binaural stethoscope in 1851. We wondered whether it might be possible to develop a disposable index finger-based system to assess anorectal function, a system we have termed “digital manometry” (DM) [10–13]. This would include not only measures of the pressure but also muscle coordination.

We tested the primary hypothesis that in three different activities, rest, squeeze, and simulated defecation, DM pressure readings are equivalent to those measured with a HR-ARM in a sample of healthy subjects, CC and FI patients. Using DM only, we also tested the secondary hypothesis that the change in the myoelectric activities of the puborectalis muscle (PR) was proportional to change in AS maximum pressure recorded by the DM device. Finally, we tested the tertiary hypothesis that there was no difference in comfort reported by individuals undergoing DM and HR-ARM.

## 2. Methods

### 2.1. Experimental design

This was a single center, cross-sectional, observational cohort study of AS pressures in three different physical activities: at rest, during maximum squeeze, and during simulated defecation measured with both the DM system and a commercially available HR-ARM system. This study was approved by the institutional review board (HUM00046068) and all subjects signed a written consent form prior to the experiment.

### 2.2. Participants

We recruited 16 participants including both healthy individuals (2 males, 2 females) and patients with a diagnosis of CC (2 males, 2 females) or FI (1 male, 7 females). The median age was 61 (range: 31 – 85) years, and the mean BMI was 29.4 (SD: 5.9) kg/m^2^. The healthy volunteers were recruited by public advertisement, free of gastrointestinal symptoms, and not taking medications affecting gastrointestinal or colonic function. The CC patients fulfilled the Rome III criteria [14] for chronic functional constipation and were referred for HR-ARM to evaluate persistent constipation symptoms despite laxative therapies. Finally, FI subjects had at least one episode of accidental bowel leakage in the previous month [2,15]. All subjects were free of structural anorectal diseases and previous anorectal or colonic surgery.

All participants underwent the same procedures with the DM and HR-ARM systems. One experienced operator (WDC) used the DM and the other (JRB) the HR-ARM device. The operators were blinded as to the other’s results. The order of performing the assessment with two measurement methods was randomized for each subject to minimize possible learning effects.

### 2.3. Apparatus

#### 2.3.1 Digital Manometry

DM consisted of a disposable, instrumented glove and a reusable wrist-mounted determining unit (WDU) (Fig 1). The glove had three miniature (1 mm × 1 mm × 0.6 mm) piezo-resistive micro-electromechanical pressure sensors (P1602 NovaSensor [100 kPa], Amphenol, CA) mounted on a 300 µm-thick custom flexible printed circuit board (FPCB) made from biocompatible polyimide substrate [16]. Each sensor was covered with a smooth uniform layer of biocompatible silicone elastomer (MED2-4220, Nusil, CA). Afterwards, in a clean environment, the FPCB was adhered to the outside of the index digit of a standard non-latex surgical glove using biocompatible silicone adhesive (MED-1011, Nusil, CA). The location of pressure sensors of the FPCB were on the tip of the distal phalange over where the operator’s nail would be, the medial aspect of the middle phalanx, and the lateral interphalangeal joint, respectively. This arrangement enables measuring intrarectal and AS pressures during the detailed digital rectal examination (see Fig 2). The base sensors (P_2_ and P_3_ shown in Fig 1) of the DM device were designed to be kept within the AS maximum pressure zone. Additionally, the FPCB had two pairs of gold-plated myoelectric electrodes, each spaced 1 mm apart over the index fingertip pad and on the aspect of the middle segment of the index finger, to measure the myoelectric activity of PR and AS muscles, respectively. The entire circuit board was covered by an extra protective layer provided by a silicone rubber finger cot with openings to expose only the myoelectric electrodes (Fig 1).

**Fig 1.**
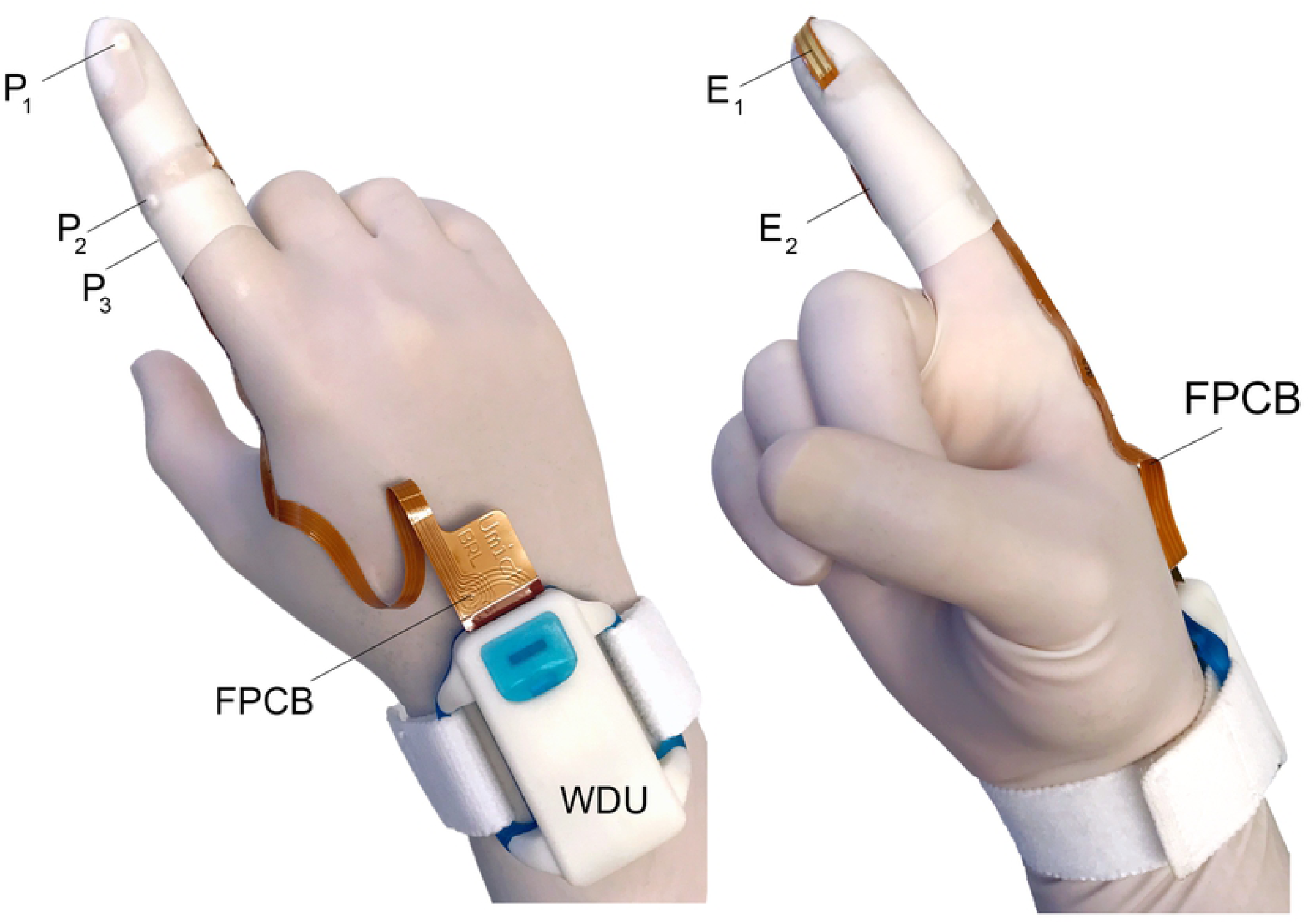
Wearable DM system. Two views of the DM disposable instrumented glove along with the reusable wrist-mounted determining unit (WDU). (Left) The three pressure sensors, P_1-3_, are mounted on the flexible printed circuit board (FPCB) that wraps around the index finger under a finger cot. (Right) The two myoelectric electrodes, E_1-2_, also connected to the FPCB, are shown adhered to the surface of the finger cot. WDU amplifies and transmit signals from the pressure sensors and myoelectric electrodes wirelessly to a nearby computer for display purposes. The patient myoelectric ground electrode and cable that connect to the WDU are not shown for simplicity.

**Fig 2.**
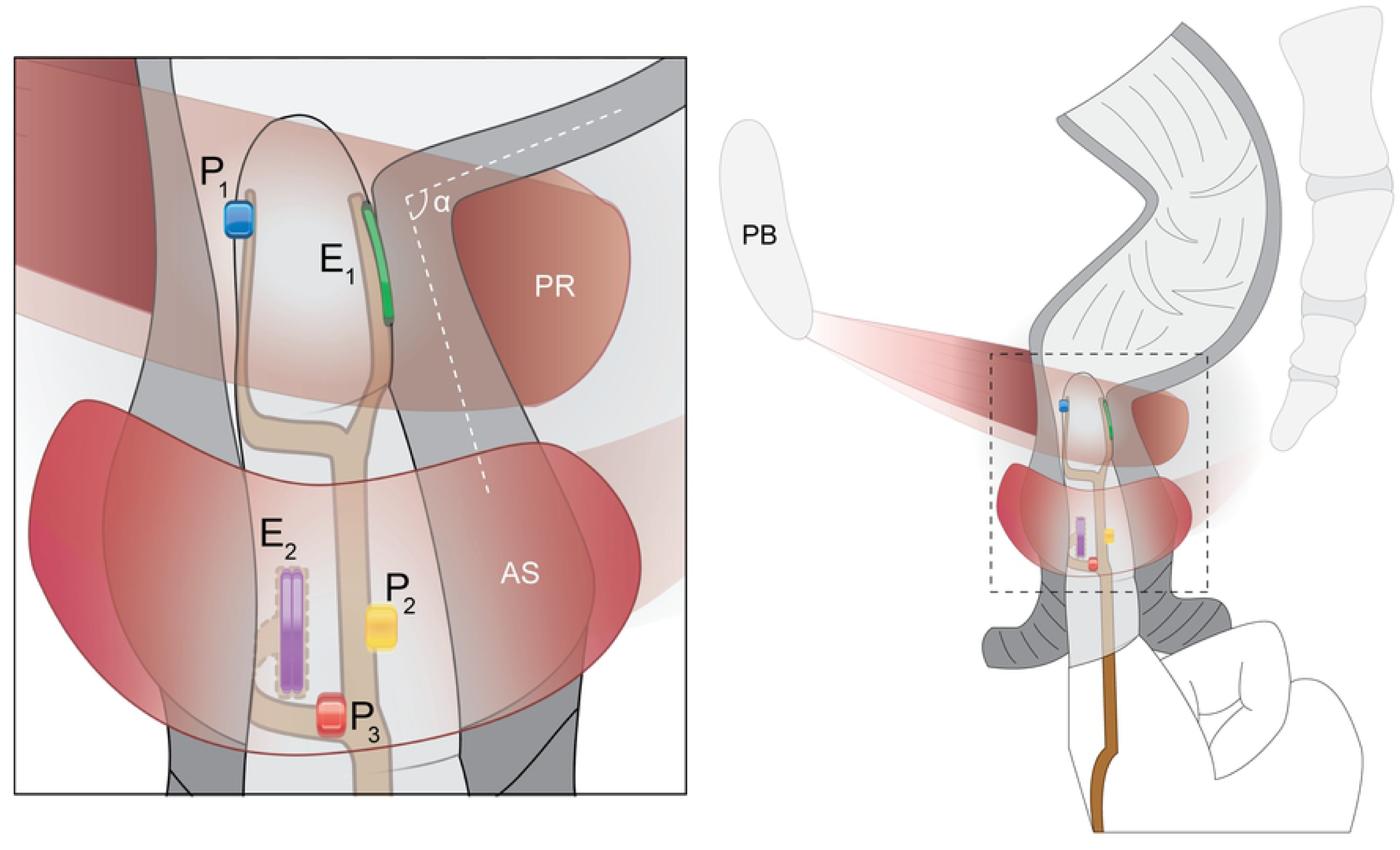
DM instrumented glove. A schematic illustration showing a left lateral view of the DM disposable glove during an anorectal canal assessment. Single pressure sensors are located over the fingernail (P_1_), middle phalanx (P_2_), and the proximal interphalangeal joint (P_3_), as well as two bipolar gold plated myoelectric activity electrodes (E_1_ and E_2_) for the puborectalis (PR) and anal sphincter (AS) muscles. The inset at left shows how the printed circuit board wraps around the index finger. α is the anorectal angle. Note that the ipsilateral PR muscle is shown as being transparent to permit a view of rectal canal.

A reusable wrist-mounted signal conditioning unit transmitted data from the index finger-based sensors to a computer for further analyses and display. This unit was comprised of a first order high-pass passive filter with a breakpoint frequency of 186 Hz to attenuate any DC offset and dual instrumentation amplifiers (INA2128, Texas Instruments, TX) to amplify myoelectric signals with a gain of 10,000. The myoelectric signal was hardware rectified and filtered (see S1 Text). Finally, we used an over-the-counter electrocardiogram electrode over the greater trochanter as the ground reference. Computers recorded pressures from the anorectal complex as well as the myoelectric activity of AS muscles at 60 Hz. The pressure and myoelectric signals were processed using a center point moving window to calculate the average of 20 data points.

#### 2.3.2 High-Resolution Manometry

A Sandhill Scientific® HR-ARM system (Denver, CO, USA) was used with a 4 mm diameter catheter (UNI-ANO-M0138) having five sets of 4 radially- and orthogonally-arranged pressure sensors spaced 1 cm apart. A single sensor on the catheter was located away from the array and exposed to the atmospheric pressure outside the rectum as a datum. The catheter had two single sensors at the distal end, one of them being inside an inflatable balloon (Fig 3).

**Fig 3.**
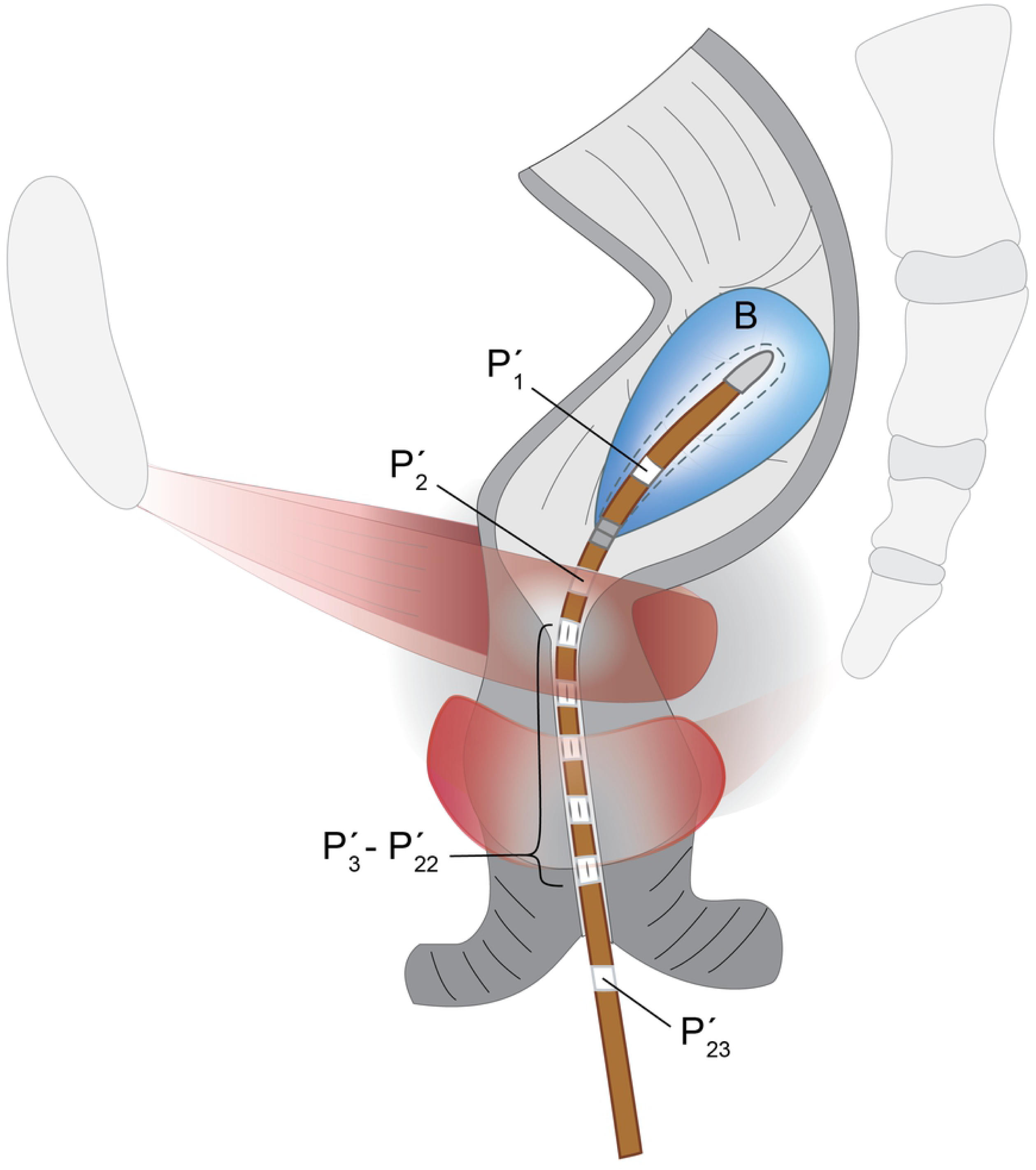
HR-ARM catheter. Illustration of the HR-ARM 4 mm–diameter catheter inside the anal canal during the anorectal manometry procedure. Also shown are, relative to the patient, the proximal 60 (cm^3^) inflated balloon (B) with its single pressure sensor inside (P’_1_), a single pressure sensor outside the balloon (P’_2_), five sets of four radially and orthogonally arranged pressure sensors (P’_3_-P’_22_), and the external pressure reference pressure sensor (P’_23_) distally.

### 2.4 Study procedures

During the clinical visit, the examiner first checked the anal canal to ensure it was free from scars and evidence of bleeding. Each participant was asked to lie in the left lateral decubitus position with both legs flexed. The reusable HR-ARM catheter was sterilized and recalibrated prior to each study. Upon insertion of either the lubricated DM probe or HR-ARM catheter, the examiners asked each subject to relax their anorectal muscles for at least one minute to measure the average resting pressures. Afterwards, participants were asked to maximally squeeze the AS muscle for 5 seconds as if they were trying to prevent accidental bowel leakage. Once the subject’s AS pressure returned to the baseline, they repeated the same maneuver with a subsequent rest period at least twice. Finally, participants were asked to simulate defecation by expelling the digit of the examiner wearing the DM instrumented glove or the HR-ARM catheter. Pressure signals were recorded continuously throughout the entire session and available to the clinician as real time visual feedback. In the case of DM, myoelectric activities were recorded simultaneously. Subjects were asked to press an event marker button at the beginning and end of each trial of each activity that they were asked to perform (Fig 4).

**Fig 4.**
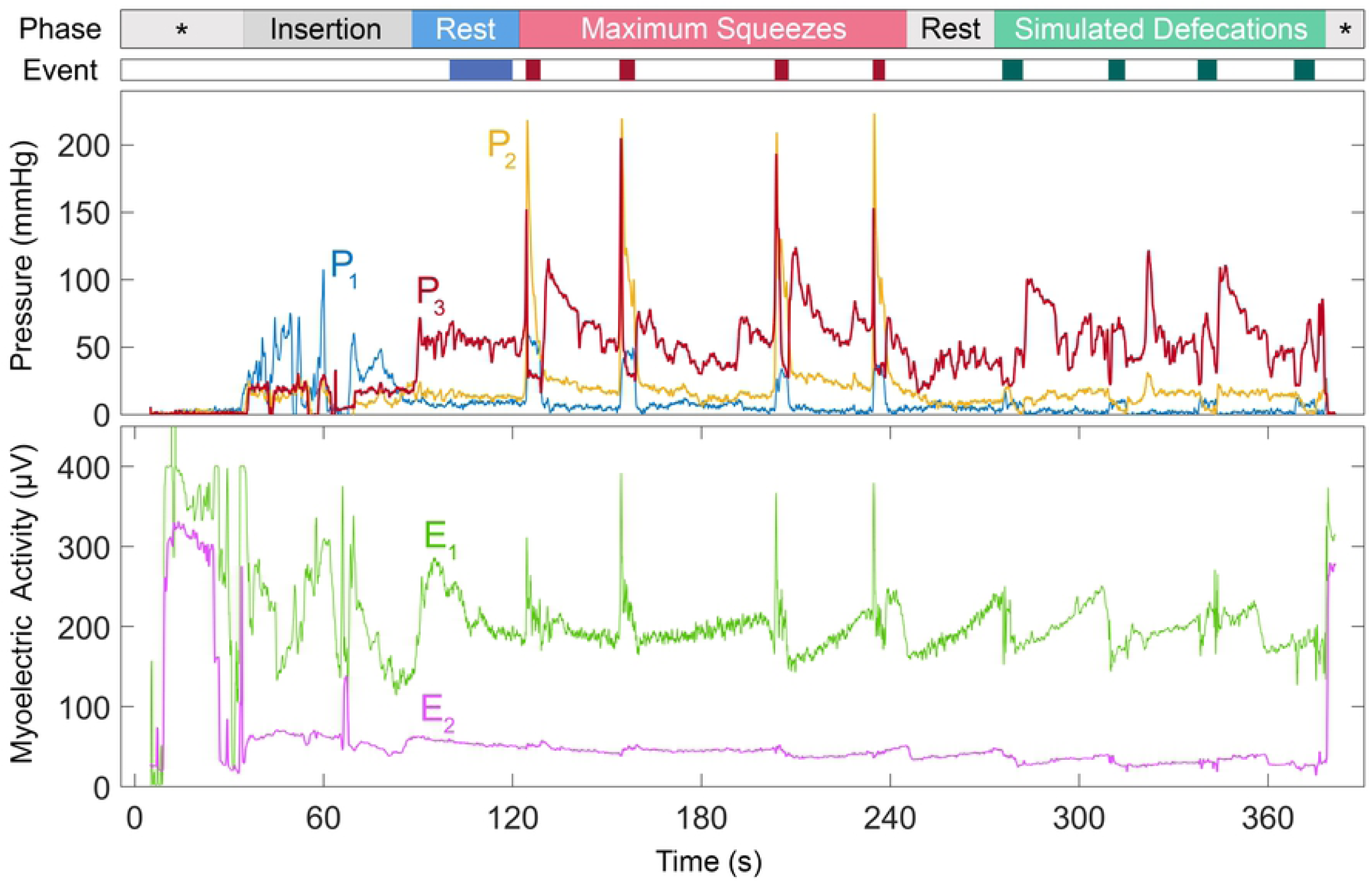
Sample recordings of the DM device. Sample simultaneous AS and rectal pressures (P1-P3) with myoelectric activities (E1 and E2) of the anorectal muscles recorded from a healthy male (subject #3). The first bar (“Phase”) along the top of the illustration shows the exam phases. The first asterisk denotes the period before the sensors are inserted into the body. Baseline activity was recorded with the patient resting (“Rest”). The patient was then asked to maximally contract their AS muscles four times interspersed by rest periods. After another rest (“Rest”), the patient was asked to simulate defecation four times (“Simulated Defecations”). The last asterisk denotes the device being removed from the body and outside the body. The second color-coded (“Event”) bar shows the intervals when the patient pressed the hand-held event marker during which measurements were made. (Note that the floating myoelectric activity signal prior to insertion of the probe at t = 35 s).

Once both procedures were completed, subjects were asked to complete a survey on the comfort level of each system. A 10-point visual analog discomfort scale was used with ‘1’ labeled ‘unbearable’, ‘5’ being ‘bearable’, and ‘10’ representing ‘very acceptable’ on three factors: smoothness, shape, and size of the probes. The ‘overall comfort level’ as well as the ‘duration’ of each method were also rated on the same scale. Finally, in the survey, subjects were asked to describe their opinion or experience during studies and whether DM could be improved in any way.

### 2.4. Statistical analyses

To test the primary hypothesis the Bland-Altman [17] limits of agreement (LoA) method was used to compare pressure readings for the rest, squeeze, and simulated defecation procedures (paired differences = DM - HR-ARM). In the field of gastroenterology, pressures are either measured in mmHg or cmH_2_O; we have chosen the former (for more familiar with SI units, 1 mmHg is equivalent to 133.3 Pa). For the rest values, mean (SD) were calculated from at least 10 seconds of DM data and mean values taken from HR-ARM output. The maximum pressure during squeeze was calculated by taking the peak value in the interval between the event markers. For each subject, the defecation episode was selected with the maximum pressure decrease from the rest value just prior to the effort to the minimum pressure measured during the maneuver. For the HR-ARM, since it had more sensors than DM, the mean pressure value from each set of four radially-arranged sensors was first calculated for each 5-second interval. Then, the maximum value was found along the catheter to represent the high pressure zone in the AS.

We calculated the absolute and relative intraobserver errors [18,19] for three consecutive maximum squeeze pressures by each subject for only DM, since the reproducibility of measurements made with the HR-ARM device have already been documented [20].

To test the secondary hypothesis, least squares regression was used to find the correlation between the changes in the myoelectric activity of the PR muscle and AS pressure change in maximum squeeze maneuver.

Finally, to compare the comfort level of both systems a two-sided Mann-Whitney U-test was used with a p level of 0.05 to compare the DM and HD-ARM ratings for comfort, smoothness, size, shape, and duration.

## 3. Results

The data support the primary hypothesis in that the LoA showed acceptable DM performance for measuring AS pressure at rest (Fig 5A) and during simulated defecation (Fig 5B). For the rectal pressure measurement, we noted greater variance (Fig 5C). The largest difference between the two methods was found in the measurement of AS pressure during a maximum squeeze (Fig 5D). If two outliers from the results of the maximum squeeze episodes were excluded (see Discussion), mean ± SD LoA decreased from −20 ± 172.6 to 7.2 ± 93.6 mmHg. The mean of the standard deviations of maximum pressures in all squeeze episodes across all subjects for DM and HR-ARM were 14.9 mmHg and 9.5 mmHg, respectively (Table 1).

**Table 1.** Results from Bland & Altman tests comparing DM and HR-ARM for the AS and rectal pressure recordings across all subjects.

| Maneuver | Mean Paired Difference (mmHg) | LOA (mmHg) (CI: 95%) |
| --- | --- | --- |
| AS pressure at rest | -7.1 | ±25.7 |
| AS pressure change in simulated defecation | 0.4 | ±23 |
| Rectal pressure change in simulated defecation | -37.6 | ±50.9 |
| AS pressure increment in maximum squeeze | -20.6 | ±172.6 |

**Fig 5.**
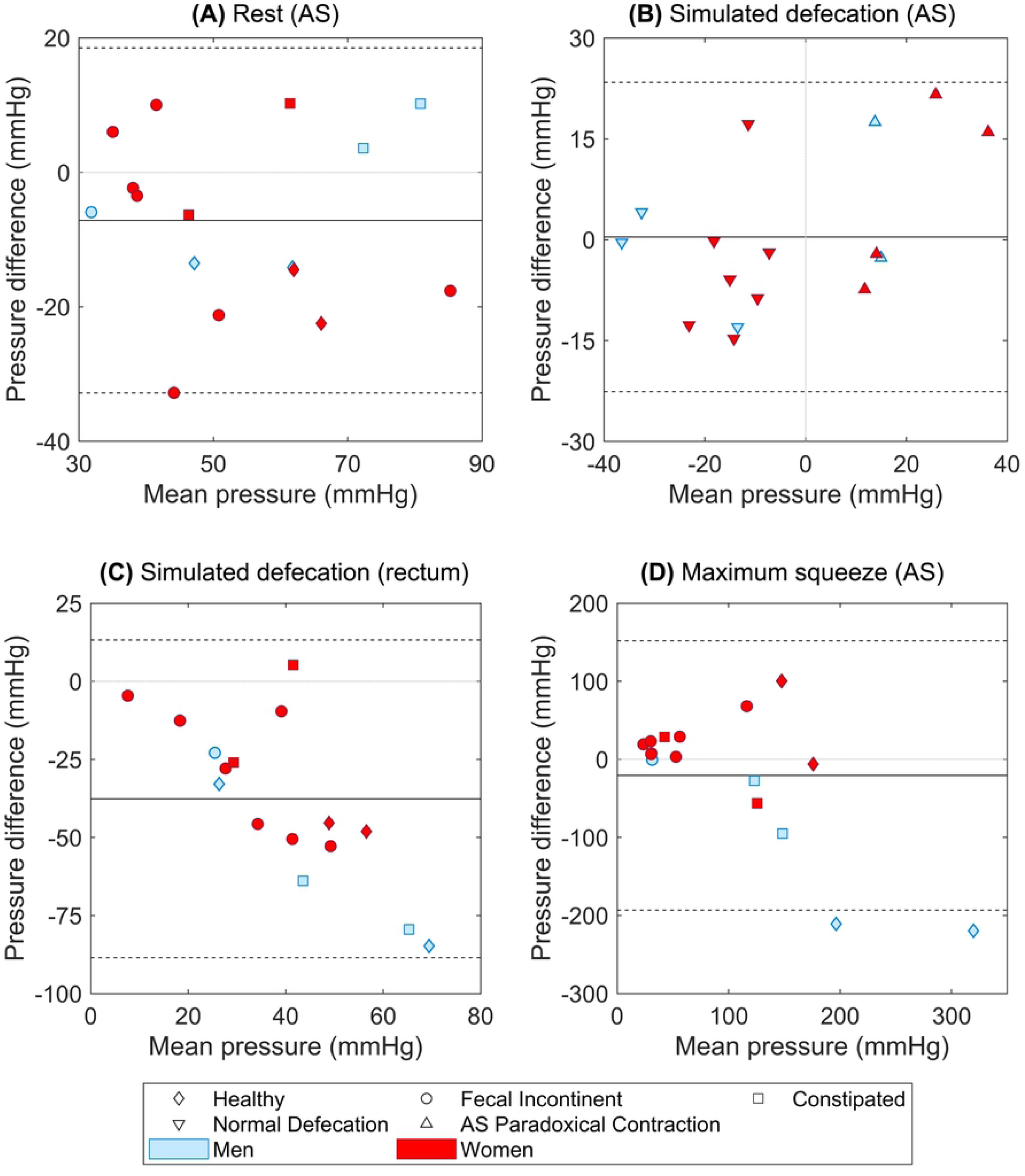
Bland & Altman comparisons between DM and HR-ARM in measuring anorectal pressures. (A) AS resting pressure, (B) AS pressure change during simulated defecation, (C) rectal pressure change during simulated defecation, and (D) AS pressure change during maximum squeeze.

During simulated defecation, 37.5% of the subjects paradoxically contracted their AS by the predicate HR-ARM. There was complete agreement between DM and HR-ARM in distinguishing paradoxical AS contraction during simulated defecation Fig 5B). The samples on the left side of the plot, show examples of subjects with normal AS relaxation during simulated defecation (i.e., negative AS pressure change), whereas the points on the right side of the plot represent subjects who paradoxically contracted their AS (i.e., positive AS pressure change).

The mean (SD) absolute and relative intraobserver variability of DM were 12 (21.2) and 14% (21%) with ICC of 0.94.

For the DM device, the slope and intercept of the regression line (Fig 6) between the percent change in the PR myoelectric activity and increment in AS squeeze pressure was found to be 0.6 µV/(µV.mmHg) and 25.5 µV/µV, respectively (R^2^ = 0.4). Therefore, the change in myoelectric activity of the PR muscle was proportional to the change in AS maximum squeeze pressure, supporting the secondary hypothesis.

**Fig 6.**
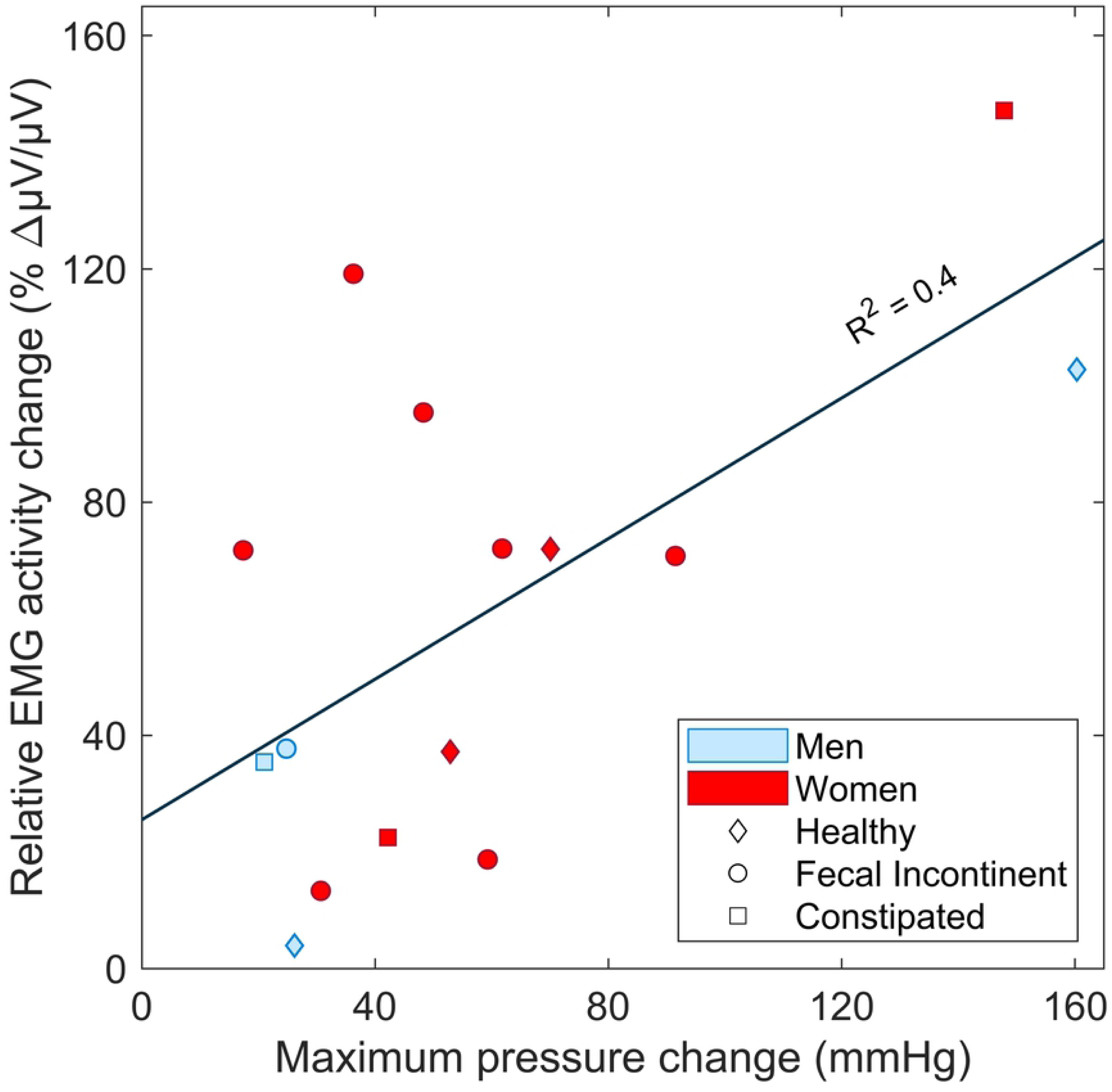
Maximum change in PR myoelectric activity vs AS pressure change in simulated defecation across all patients. Trials were selected which showed the largest change in peak PR myoelectric activity compared with the maximum decrease in the AS pressure during that defecation maneuver in 15 of 16 subjects (no myoelectric activity was recorded in the first subject due to equipment malfunction). Note that the patients with dyssynergic defecation could not decrease AS pressure (see Fig 4B).

Finally, the results from surveys (Table 2) showed that the difference between comfort levels of both systems were not significant (Table 3).

**Table 2.**
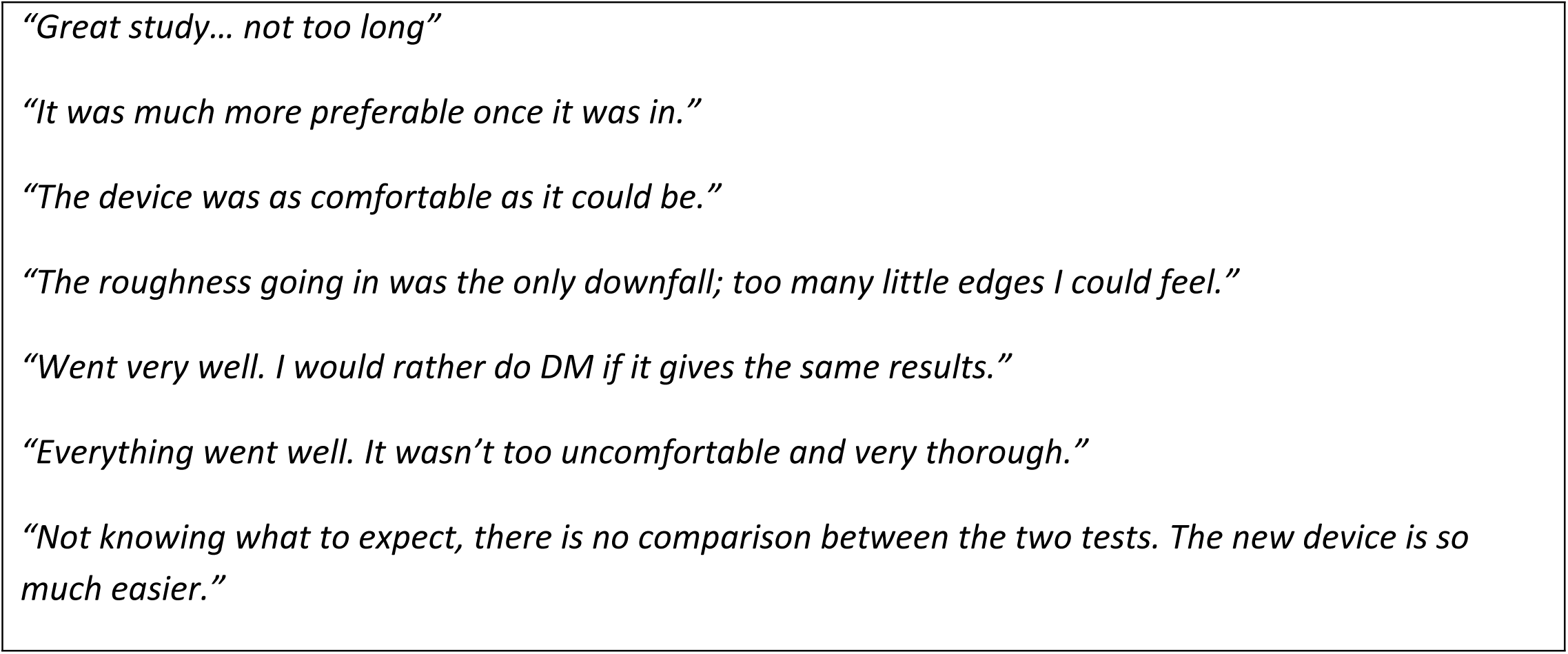
Examples of the participants’ responses to the survey question on DM system comfort.

**Table 3.** Comparison of DM and HR-ARM subjective post-hoc test assessment levels on a visual analog scale from 1 to 10, with ‘1’ labeled ‘unbearable’ and ‘10’ representing ‘very acceptable’. ‘Min’ denotes minimum and ‘max’ denotes maximum.

| Factor | DM |  |  | HR-ARM |  |  | P-Value |
| --- | --- | --- | --- | --- | --- | --- | --- |
|  | Mean (SD) | Min | Max | Mean (SD) | Min | Max |  |
| Comfort | 9.3 (1.0) | 7 | 10 | 9.1 (1.5) | 5 | 10 | 1 |
| Smoothness | 9.6 (0.8) | 8 | 10 | 9.2 (1.6) | 4 | 10 | 0.56 |
| Shape | 9.6 (1.0) | 7 | 10 | 9.3 (1.6) | 5 | 10 | 0.67 |
| Size | 9.3 (1.1) | 7 | 10 | 9.3 (1.8) | 4 | 10 | 0.68 |
| Duration | 9.9 (0.5) | 8 | 10 | 9.3 (1.7) | 4 | 10 | 0.35 |

## 4. Discussion

Although physicians have worn wearables in the form of stethoscopes and loupes to augment their sensory acuity for over a century, the miniaturization of sensors and electronics is providing new opportunities for developing wearables for clinicians to use to evaluate patients quickly and easily. This paper describes a novel wearable that allows providers to better understand the etiology of chronic constipation and fecal incontinence at the point of service. If validated by others, digital manometry could, for the first time, offer providers in low-resource settings the ability to gain information that would otherwise require referral to a tertiary center for comprehensive anorectal function testing. Further, we describe an early phase prototype. It is certainly conceivable that the instrumented glove and wrist-mounted determining unit can be linked to a smart phone via Bluetooth. In this paper, we have chosen to report pressure in units of mmHg because gastroenterologists interested in this paper are familiar with typical ranges for those units rather than the SI units standard in engineering and science.

Our primary hypothesis was supported in that the DM provided accurate pressure readings compared to the gold standard HR-ARM system. It did so at a hardware cost that is an order of magnitude less than HR-ARM (Table 4). More specifically, DM was equivalent to the gold standard HR-ARM system in measuring the AS pressure during rest and simulated defecation episodes.

**Table 4.** Estimated cost comparison between DM and HR-ARM methods.

|  | DM | HR-ARM |
| --- | --- | --- |
| Overall system cost | \$10,000 | \$122,000 |
| Overall hardware cost | \$1,500 | \$116,000 |
| Probe cost | \$10 | \$6,000 <sup>a</sup> |
| Sterilization cost <sup>b</sup> | \$0 <sup>c</sup> | \$20 |
| Hardware cost per exam | \$10 | \$50 |

<sup>a</sup> 200 usages per catheter.
<sup>b</sup> per session.
<sup>c</sup> disposable instrumented glove.

In the maximum squeeze maneuver, DM recorded systematically lower average AS pressures compared to HR-ARM. Most of that difference was caused by the results from two men (Fig 5D - blue outliers). Interestingly, the literature reports higher pressure readings of HR-ARM compared to the standard conventional manometry [21–25], especially in men compared to women [26]. Indeed, recent papers on HR-ARM have suggested that new standards and limits are needed for HR-ARM based on the catheter type and size to address systematic differences in pressure measurements [25,27].

We believe that there are three main reasons why there was a difference between recorded AS pressures using DM and HR-ARM. The first is the hypersensitivity of HR-ARM in squeeze test [24,25,27], which could be due to the relatively high bending stiffness of its probe. This would cause the HR-ARM probe to measure high tissue contact stress (force per unit area) in the vicinity of its tip as well as where the shaft is bent the most over the anorectal angle, rather than just measuring fluid pressure within the AS or rectum [28]. On the other hand, although the sensors in the DM probe are supported by index finger tissues, the finger can be purposely relaxed to adopt the shape of the anorectal angle thereby reducing tissue contact stress. We believe the reason that DM measured smaller pressure change than HR-ARM in three activities (Table 1) was because we detuned the pressure sensitivity of the sensor by thickening the silicone layer over the sensor die in order to improve its sensitivity to lower pressures. Whereas, the Bland-Altman analysis treats the HR-ARM as the gold standard measurement, we conclude that our results corroborate the literature suggesting HR-ARM has its own measurement biases caused by its having a catheter that is stiffer than the anorectum in bending.

The test of the secondary hypothesis showed that the change in the relative myoelectric activity of the PR muscle measured by the electrode at the distal end of the examining finger was proportional to the pressure change in the AS pressure during maximum squeeze maneuver (Fig 6). Unfortunately, the examining finger tended to often be partially expelled during attempted defecation making the PR muscle myoelectric measurements unreliable.

The combined pressure and myoelectric activity data provided by DM enables the identification of dyssynergic defecation caused by poor coordination of the abdominal wall and anorectal muscles. A primer of DM and HR-ARM results indicative of dyssynergic defecation can be found in Table 5.

**Table 5.** AS pressure change from rest to activity measured with DM and HR-ARM; positive changes represent dyssynergic defecation.

| Subject | AS pressure change (mmHg) |  | Defecation |
| --- | --- | --- | --- |
|  | DM | HR-ARM |  |
| 1 | 8 | 15.4 | Dyssynergic |
| 2 | -30.5 | -34.6 | Normal |
| 3 | -36.7 | -36.3 | Normal |
| 4 | -8.2 | -6.3 | Normal |
| 5 | -13.9 | -5.2 | Normal |
| 6 | -18 | -12.1 | Normal |
| 7 | -21.6 | -6.9 | Normal |
| 8 | 13 | 15.1 | Dyssynergic |
| 9 | -2.8 | -20 | Normal |
| 10 | 44.2 | 28.2 | Dyssynergic |
| 11 | -20 | -7 | Normal |
| 12 | 22.5 | 5 | Dyssynergic |
| 13 | -18.3 | -18.1 | Normal |
| 14 | -29.5 | -16.8 | Normal |
| 15 | 36.6 | 15 | Dyssynergic |
| 16 | 13.6 | 16.3 | Dyssynergic |

The comfort surveys from the participants suggest that DM provides comparable levels of comfort and usability even though the HR-ARM probe was a quarter of the diameter of the DM covered index finger (Table 3). The lowest score received by DM on any of the five variables was a 7 out of 10, whereas HR-ARM was a 4 out of 10. The slightly higher average score of DM on the smoothness could be due to the silicone cover layer over the instrumentations of the glove.

A major limitation of this feasibility study was the small sample size in each group. Since this was a feasibility study, we were interested in studying healthy subjects as controls and two common patient types. The data showed that it is feasible to mount pressure sensors on the index finger, obtain meaningful measurement of anorectal pressures, and observe paradoxical contraction of AS muscles during simulated defecation. More subjects in prospective DM studies are needed to corroborate and extend the present results and define normal pressure ranges [23,29,30].

A second limitation was the use of one DM and one HR-ARM observer and although we could calculate the intraobserver error, we could not calculate the interobserver error for either modality. The use of the single observer also meant that the effect of different index finger diameters and lengths on the DM measured pressure recordings could not be examined. Although the breadth of the PIP joint of this examiner was 19 mm, pressure recordings of AS muscle using a rigid 20 mm diameter probe can average 37 mmHg and 50 mmHg higher than with a rigid 4 mm probe [31] during rest and maximum squeeze. The fact that on average the DM readings were 7.1 mmHg less relative to HR-ARM suggests that the compliance and shape change of the finger in the glove can compensate for the effect of the finger having a larger diameter than the 4 mm probe. But one can anticipate a range of clinician index finger mean diameters ranging from 1.8 mm to 2.3 mm [32], so future studies should examine the effect of examiner digit finger diameter and anthropometry on measured anorectal pressures. It may be possible to create a look up table or nomogram to interpret the measured pressures.

A third limitation was the axial movement of either probe away from the AS high pressure zone during the maximum squeeze and simulated defecation maneuverers. In HR-ARM, the recordings show that the region of high pressure moves about 1 cm proximally on the catheter during the maximum squeeze. Similarly, in DM, although the examiner tried to minimize the probe displacement from the high AS pressure zone, some sensor movements relative to high pressure zone was inevitable because the examiner can never exactly match the movement of the patient’s anorectal complex. This is reflected in the finding that 1 cm displacement from high pressure zone in the AS can result in a 30% reduction in pressure reading [31]. Therefore, both systems were susceptible to movement artifact, but the advantage of the HD-ARM is that it provides redundancy by multiple measurements over a longer length of the probe. Such redundancy in the number and arrangement of sensors can be incorporated in future DM systems.

Since DM probes are disposable, they eliminate the risk of patient-to-patient transmission of infection compared with HR-ARM. Even though the HR-ARM system provides detailed spatial pressure recordings throughout the AS complex, it can be cost prohibitive for many patients and health care providers. Therefore, if validated by others, DM could provide an inexpensive anorectal screening modality for a provider to determine next best steps for a patient with dyssynergic defecation or fecal incontinence. For example, if DM is abnormal, a provider might decide to refer the patient to a tertiary medical center or perhaps directly to biofeedback and pelvic floor physical therapy. Clearly, further studies will also be needed to determine how best to use the information provided by DM.

## 5. Conclusions

1. DM provided comparable recordings of resting AS pressure, as well as the important changes in anorectal pressures during prescribed activities, as HR-ARM. Further validation requires larger sample sizes, and interobserver differences should be examined.
2. DM myoelectric data can help identify paradoxical contraction of the AS and PR muscles suggestive of dyssynergic defecation.
3. The two systems had similar comfort levels despite different probe diameters.

## Author contributions

**Conceptualization:** Ali Attari, William Chey, James Ashton-Miller

**Data Curation:** Ali Attari

**Formal Analysis:** Ali Attari

**Funding Acquisition:** William Chey, James Ashton-Miller

**Investigation:** Ali Attari, William Chey, Jason Baker

**Methodology:** Ali Attari, William Chey, Jason Baker, James Ashton-Miller

**Project Administration:** James Ashton-Miller, William Chey.

**Resources:** William Chey, James Ashton-Miller

**Software:** Ali Attari

**Supervision:** William Chey, James Ashton-Miller

**Validation:** William Chey, James Ashton-Miller

**Visualization:** Ali Attari

**Writing – Original Draft Preparation:** Ali Attari, James Ashton-Miller

**Writing – Review & Editing:** Ali Attari, William Chey, Jason Baker, James Ashton-Miller

## References

1. Carrington E V., Scott SM, Bharucha A, Mion F, Remes-Troche JM, Malcolm A, et al. Advances in the evaluation of anorectal function. Nat Rev Gastroenterol Hepatol [Internet]. 2018 May 11;15(5):309–23. Available from: http://dx.doi.org/10.1038/nrgastro.2018.27

2. Whitehead WE, Borrud L, Goode PS, Meikle S, Mueller ER, Tuteja A, et al. Fecal Incontinence in US Adults: Epidemiology and Risk Factors. Gastroenterology [Internet]. 2009 Aug;137(2):512-517.e2. Available from: https://linkinghub.elsevier.com/retrieve/pii/S0016508509007215

3. Johanson JF, Lafferty J. Epidemiology of fecal incontinence: the silent affliction. Am J Gastroenterol [Internet]. 1996 Jan [cited 2017 Mar 9];91(1):33–6. Available from: http://www.ncbi.nlm.nih.gov/pubmed/8561140

4. Higgins PDR, Johanson JF. Epidemiology of Constipation in North America: A Systematic Review. Am J Gastroenterol [Internet]. 2004 Apr;99(4):750–9. Available from: http://www.nature.com/doifinder/10.1111/j.1572-0241.2004.04114.x

5. Dinning PG, Carrington E V., Scott SM. The use of colonic and anorectal high-resolution manometry and its place in clinical work and in research. Neurogastroenterol Motil [Internet]. 2015 Dec;27(12):1693–708. Available from: http://doi.wiley.com/10.1111/nmo.12632

6. Dinning PG, Carrington E V., Scott SM. Colonic and anorectal motility testing in the high-resolution era. Curr Opin Gastroenterol [Internet]. 2016 Jan;32(1):44–8. Available from: http://content.wkhealth.com/linkback/openurl?sid=WKPTLP:landingpage&an=00001574-201601000-00010

7. Portalatin M, Winstead N. Medical Management of Constipation. Clin Colon Rectal Surg [Internet]. 2012 Mar 23;25(01):012–9. Available from: http://www.thieme-connect.de/DOI/DOI?10.1055/s-0032-1301754

8. Norton C, Emmanuel A, Stevens N, Scott SM, Grossi U, Bannister S, et al. Habit training versus habit training with direct visual biofeedback in adults with chronic constipation: study protocol for a randomised controlled trial. Trials [Internet]. 2017 Dec 24;18(1):139. Available from: http://trialsjournal.biomedcentral.com/articles/10.1186/s13063-017-1880-0

9. Locke GR, Pemberton JH, Phillips SF. American Gastroenterological Association medical position statement: Guidelines on constipation. Gastroenterology [Internet]. 2000 Dec;119(6):1761–6. Available from: https://linkinghub.elsevier.com/retrieve/pii/S0016508500700230

10. Chey WD, Ashton-Miller JA, Spiegel BMR. Digital manometry finger-mountable sensor device [Internet]. USA; WO2013090681 A3, 2012 [cited 2016 Mar 2]. Available from: https://www-google-com.proxy.lib.umich.edu/patents/US20130158365

11. Chey WD, Baker J, Attari A, Spiegel BM, Ashton-Miller JA. A Glove-Based, Disposable, Point-of-Service Device Which Allows Detailed Physiological Assessment of the Anorectum: A Proof of Concept Study in Healthy Volunteers. Gastroenterology [Internet]. 2014 May [cited 2016 Mar 2];146(5):S–720. Available from: https://www.infona.pl//resource/bwmeta1.element.elsevier-1209e0d1-9b3a-34bf-bcfd-78c67e7a289b

12. Chey WD, Baker J, Attari A, Ashton-Miller JA. A Glove-Based, Disposable, Point-of-Service Device Which Allows Physiological Assessment of the Anorectum: Concordance With Anorectal Manometry in Chronic Constipation Patients. Gastroenterology [Internet]. 2015 Apr 1 [cited 2016 Mar 2];148(4):S–178. Available from: http://www.gastrojournal.org/article/S0016508515305941/fulltext

13. Attari A, Chey WD, Baker J, Ashton-Miller JA. Development of a Disposable Point-of-Service Digital Manometry Device to Assess Anorectal Function. In: American Society of Biomechanics [Internet]. Boulder; 2017. p. 645–6. Available from: http://asbweb.org/conferences/2017/abstracts/ASB2017_Abstracts.pdf

14. Longstreth GF, Thompson WG, Chey WD, Houghton LA, Mearin F, Spiller RC. Functional Bowel Disorders. Gastroenterology [Internet]. 2006 Apr;130(5):1480–91. Available from: https://linkinghub.elsevier.com/retrieve/pii/S0016508506005129

15. Jorge JM, Wexner SD. Etiology and management of fecal incontinence. Dis Colon Rectum [Internet]. 1993 Jan;36(1):77–97. Available from: http://www.ncbi.nlm.nih.gov/pubmed/8416784

16. Richardson RR, Miller JA, Reichert WM. Polyimides as biomaterials: preliminary biocompatibility testing. Biomaterials [Internet]. 1993 Jan;14(8):627–35. Available from: https://linkinghub.elsevier.com/retrieve/pii/0142961293901833

17. Bland JM, Altman DG. Statistical methods for assessing agreement between two methods of clinical measurement. Lancet (London, England) [Internet]. 1986 Feb 8 [cited 2017 Mar 8];1(8476):307–10. Available from: http://www.ncbi.nlm.nih.gov/pubmed/2868172

18. Popovic ZB, Thomas JD. Assessing observer variability: a user’s guide. Cardiovasc Diagn Ther [Internet]. 2017 Jun;7(3):317–24. Available from: http://cdt.amegroups.com/article/view/14271/15027

19. McGraw KO, Wong SP. Forming inferences about some intraclass correlation coefficients. Psychol Methods [Internet]. 1996 Mar;1(1):30–46. Available from: http://doi.apa.org/getdoi.cfm?doi=10.1037/1082-989X.1.1.30

20. Chakraborty S, Feuerhak KJ, Zinsmeister AR, Bharucha AE. Reproducibility of high-definition (3D) manometry and its agreement with high-resolution (2D) manometry in women with fecal incontinence. Neurogastroenterol Motil [Internet]. 2017 Mar;29(3):e12950. Available from: http://doi.wiley.com/10.1111/nmo.12950

21. Vitton V, Ben Hadj Amor W, Baumstarck K, Grimaud JC, Bouvier M. Water-perfused manometry vs three-dimensional high-resolution manometry: a comparative study on a large patient population with anorectal disorders. Color Dis [Internet]. 2013 Dec;15(12):e726–31. Available from: http://doi.wiley.com/10.1111/codi.12397

22. Lee HJ, Jung KW, Han S, Kim JW, Park S-K, Yoon IJ, et al. Normal values for high-resolution anorectal manometry/topography in a healthy Korean population and the effects of gender and body mass index. Neurogastroenterol Motil [Internet]. 2014 Apr;26(4):529–37. Available from: http://www.ncbi.nlm.nih.gov/pubmed/25418939

23. Carrington E V., Brokjær A, Craven H, Zarate N, Horrocks EJ, Palit S, et al. Traditional measures of normal anal sphincter function using high-resolution anorectal manometry (HRAM) in 115 healthy volunteers. Neurogastroenterol Motil [Internet]. 2014 May;26(5):625–35. Available from: http://doi.wiley.com/10.1111/nmo.12307

24. Carrington E V., Knowles CH, Grossi U, Scott SM. High-resolution Anorectal Manometry Measures Are More Accurate Than Conventional Measures in Detecting Anal Hypocontractility in Women With Fecal Incontinence. Clin Gastroenterol Hepatol [Internet]. 2019 Feb;17(3):477-485.e9. Available from: https://doi.org/10.1016/j.cgh.2018.06.037

25. Gosling J, Plumb A, Taylor SA, Cohen R, Emmanuel A V. High-resolution anal manometry: Repeatability, validation, and comparison with conventional manometry. Neurogastroenterol Motil [Internet]. 2019 Jun 15;31(6):e13591. Available from: https://onlinelibrary.wiley.com/doi/abs/10.1111/nmo.13591

26. Feuerhak K, Chakraborty S, Tirumanisetty P, Bharucha AE. Sex Differences in Anorectal Pressures and Mechanisms of Defecation in Healthy People. Gastroenterology [Internet]. 2017 Apr;152(5):S318–9. Available from: http://dx.doi.org/10.1016/S0016-5085(17)31342-2

27. Rasijeff AMP, Withers M, Burke JM, Jackson W, Scott SM. High-resolution anorectal manometry: A comparison of solid-state and water-perfused catheters. Neurogastroenterol Motil [Internet]. 2017 Nov;29(11):e13124. Available from: http://doi.wiley.com/10.1111/nmo.13124

28. Simpson RR, Kennedy ML, Nguyen MH, Dinning PG, Lubowski DZ. Anal Manometry: A Comparison of Techniques. Dis Colon Rectum [Internet]. 2006 Jul;49(7):1033–8. Available from: https://insights.ovid.com/crossref?an=00003453-200649070-00012

29. Noelting J, Ratuapli SK, Bharucha AE, Harvey DM, Ravi K, Zinsmeister AR. Normal Values for High-Resolution Anorectal Manometry in Healthy Women: Effects of Age and Significance of Rectoanal Gradient. Am J Gastroenterol [Internet]. 2012 Oct;107(10):1530–6. Available from: http://insights.ovid.com/crossref?an=00000434-201210000-00015

30. Ratuapli SK, Bharucha AE, Noelting J, Harvey DM, Zinsmeister AR. Phenotypic Identification and Classification of Functional Defecatory Disorders Using High-Resolution Anorectal Manometry. Gastroenterology [Internet]. 2013 Feb;144(2):314-322.e2. Available from: http://dx.doi.org/10.1053/j.gastro.2012.10.049

31. Gibbons CP, Bannister JJ, Trowbridge EA, Read NW. An analysis of anal sphincter pressure and anal compliance in normal subjects. Int J Colorectal Dis [Internet]. 1986 Oct;1(4):231–7. Available from: http://www.ncbi.nlm.nih.gov/pubmed/3598317

32. Greiner TM. Hand Anthropometry of U.S. Army Personnel. 1991 Dec 1 [cited 2016 Apr 10];66–73. Available from: http://oai.dtic.mil/oai/oai?verb=getRecord&metadataPrefix=html&identifier=ADA244533

